# Low-level mitochondrial heteroplasmy modulates DNA replication, glucose metabolism and lifespan in mice

**DOI:** 10.1101/179622

**Authors:** Misa Hirose, Paul Schilf, Yask Gupta, Kim Zarse, Axel Künstner, Hauke Busch, Junping Yin, Marvin N Wright, Andreas Ziegler, Marie Vallier, Meriem Belheouane, John F Baines, Diethard Tautz, Kornelia Johann, Rebecca Oelkrug, Jens Mittag, Hendrik Lehnert, Alaa Othman, Olaf Jöhren, Markus Schwaninger, Cornelia Prehn, Jerzy Adamski, Kensuke Shima, Jan Rupp, Robert Haesler, Georg Fuellen, Rüdiger Köhling, Michael Ristow, Saleh M Ibrahim

## Abstract

Mutations in mitochondrial DNA (mtDNA) lead to heteroplasmy, i.e. the intracellular coexistence of wild-type and mutant mtDNA strands, which impact a wide spectrum of diseases but also physiological processes, including endurance exercise performance in athletes. However, the phenotypic consequences of limited levels of naturally-arising heteroplasmy have not been experimentally studied to date. We hence generated a conplastic mouse strain carrying the mitochondrial genome of a AKR/J mouse strain (B6-mt^AKR^) together with a C57BL/6J nuclear genomic background, leading to >20% heteroplasmy in the origin of light-strand DNA replication (OriL). These conplastic mice demonstrate a shorter lifespan as well as dysregulation of multiple metabolic pathways, culminating in impaired glucose metabolism, compared to wild-type C57BL/6J mice carrying lower levels of heteroplasmy. Our results indicate that physiologically relevant differences in mtDNA heteroplasmy levels at a single, functionally important site impair metabolic health and lifespan in mice.

**Highlights:**

- We identify heteroplasmy of the adenine-repeat variation (9 to 13A) in nt5172 in the origin of light-strand DNA replication (OriL) in inbred mice.
- B6-mt^AKR^ mice carry >20% 12A heteroplasmy in the OriL, while B6 mice carry only ∼ 10% heteroplasmy.
- The level of 12A heteroplasmy correlates to mtDNA copy number, glucose metabolism, and lifespan in mice.
- Given the established role of mtDNA heteroplasmy in regards to endurance exercise performance in athletes, these findings may impact our understanding of metabolism and aging in humans.

## Introduction

Mitochondria play a critical role to maintain cellular activities by generating energy in the form of adenosine triphosphate (ATP) (Wallace, 2005). Additionally, mitochondria function as a signaling platform, e.g. mitochondrial reactive oxygen species control a wide range of biological processes, including epigenetics, autophagy, and immune responses (Chandel, 2014). Since mitochondria are involved in such critical cellular activities, mitochondrial dysfunction has been linked to various degenerative and metabolic conditions (e.g. Alzheimer´s disease and diabetes), cancer, and aging in humans, and this is supported by experimental evidence (Dillin et al., 2002; Feng et al., 2001; Lin and Beal, 2006; Wang et al., 2015).

Mitochondria carry their own mitochondrial DNA (mtDNA) that encodes codes 13 OXPHOS complex genes, two ribosomal RNA genes, and 22 transfer RNA genes (Anderson et al., 1981). The mode of inheritance of mtDNA is strictly maternal, and hundreds to thousands of mtDNA copies exist in a cell (Wallace, 2005). The mutations in mtDNA have been categorized into three groups: deleterious mutations, somatic mutations and adaptive polymorphisms (Wallace, 2005). Deleterious mutations result in severe mitochondrial dysfunction and are causal for maternally-inherited mitochondrial disease such as Leber hereditary optic neuropathy (LHON)(Wallace et al., 1988a) and mitochondrial encephalomyopathy, lactic acidosis, and stroke-like episodes (MELAS)(Wallace et al., 1988b). Somatic mtDNA mutations accumulate within various tissues with age, and it was experimentally shown that increased somatic mtDNA mutations exhibit aging phenotypes in mice (Kujoth et al., 2005; Trifunovic et al., 2004). In contrast, adaptive polymorphisms may be associated with survival under different climatic conditions or nutritional availability (Wallace, 2005).

Conplastic mouse strains are a unique and powerful tool to investigate the impact of mutations in mtDNA under a wide spectrum of physiological and pathological alterations (Sharpley et al., 2012), including aging (Kauppila et al., 2016; Latorre-Pellicer et al., 2016; Hirose et al., 2016). Of note, the study by Kauppila *et al.* investigating the impact of heteroplasmy on lifespan reports that higher levels of maternally inherited *de novo* mutations/heteroplasmy lead to severe pathological consequences (Kauppila et al., 2016). Higher levels of mutations/heteroplasmy rarely occur naturally. In contrast, lower levels of maternally inherited heteroplasmy commonly exist (Li et al., 2010; Payne et al., 2013), while their phenotypic consequences are to date not experimentally studied.

We previously generated a series of conplastic mouse strains, which carry distinct and stable mutations over generations in mtDNA on a C57BL/6J nuclear genomic background (Yu et al., 2009) and provide a unique opportunity to study the impact of natural variation of mtDNA on various biological and pathological processes. Since the mtDNA of those strains was previously sequenced using Sanger sequencing, which did not allow us to accurately determine levels of heteroplasmy, we here performed next generation sequencing of the mtDNA of all of our previously constructed conplastic strains and discovered a stable, maternally inherited, and low-level heteroplasmic mutation at nt5172 in the origin of L-strand replication. Moreover, the levels of the heteroplasmy varied between C57BL/6J and C57BL/6J-mt^AKR^ strains. Using this unique resource, we studied the impact of natural low-level heteroplasmy on aging, and demonstrate its consequences including an impact on mtDNA copy number ratio and the regulation of metabolic processes, which may be causative for a shorter lifespan.

## Results

### Deep-sequencing of mtDNA prepared from B6-mt^AKR^ reveals the presence of low levels of a heteroplasmic mutation at position 5172 in the OriL

We here deep-sequenced the mtDNA of a series of conplastic strains that we previously generated (Yu et al., 2009), and identified a strain carrying low levels of heteroplasmy at position 5172 in OriL. This particular strain carries the mtDNA of AKR/J (C57BL/6J-mt^AKR/J^; B6-mt^AKR^) on a C57BL/6J (B6) background. Consistent with previously published data (Bayona-Bafaluy et al., 2003; Wanrooij et al., 2012), the adenine-repeat number varies among individuals at this position. Specifically, the majority of mtDNA (approximately 60-70%) carries eleven adenines (“11A”), while the remaining percentage exhibits either 9, 10 or more than 11 adenines (9A, 10A, 12A, 13A). B6-mt^AKR^ mice have higher levels of >11A heteroplasmy compared to B6 (**Figure 1A**, 12A and 13A; *P<0.0001* for both, two-way ANOVA). While the levels of heteroplasmy vary individually, overall B6-mt^AKR^ exhibits >20% heteroplasmy in comparison to B6, which carries approximately 10% heteroplasmy at the same position in different organs and at different ages (**Figure 1B**). Interestingly, the heteroplasmy at 5172 is also observed in free-living house mice (*Mus musculus domesticus*) caught on farms (n=215, **Figure 1C**), and the pattern of heteroplasmy resembles that of B6 (**Figure 1A**). This indicates that the patterns observed in inbred lab mice are also present in nature, where they may represent a consequence of mutation pressure and possibly be subject to natural selection.

**Figure 1:**
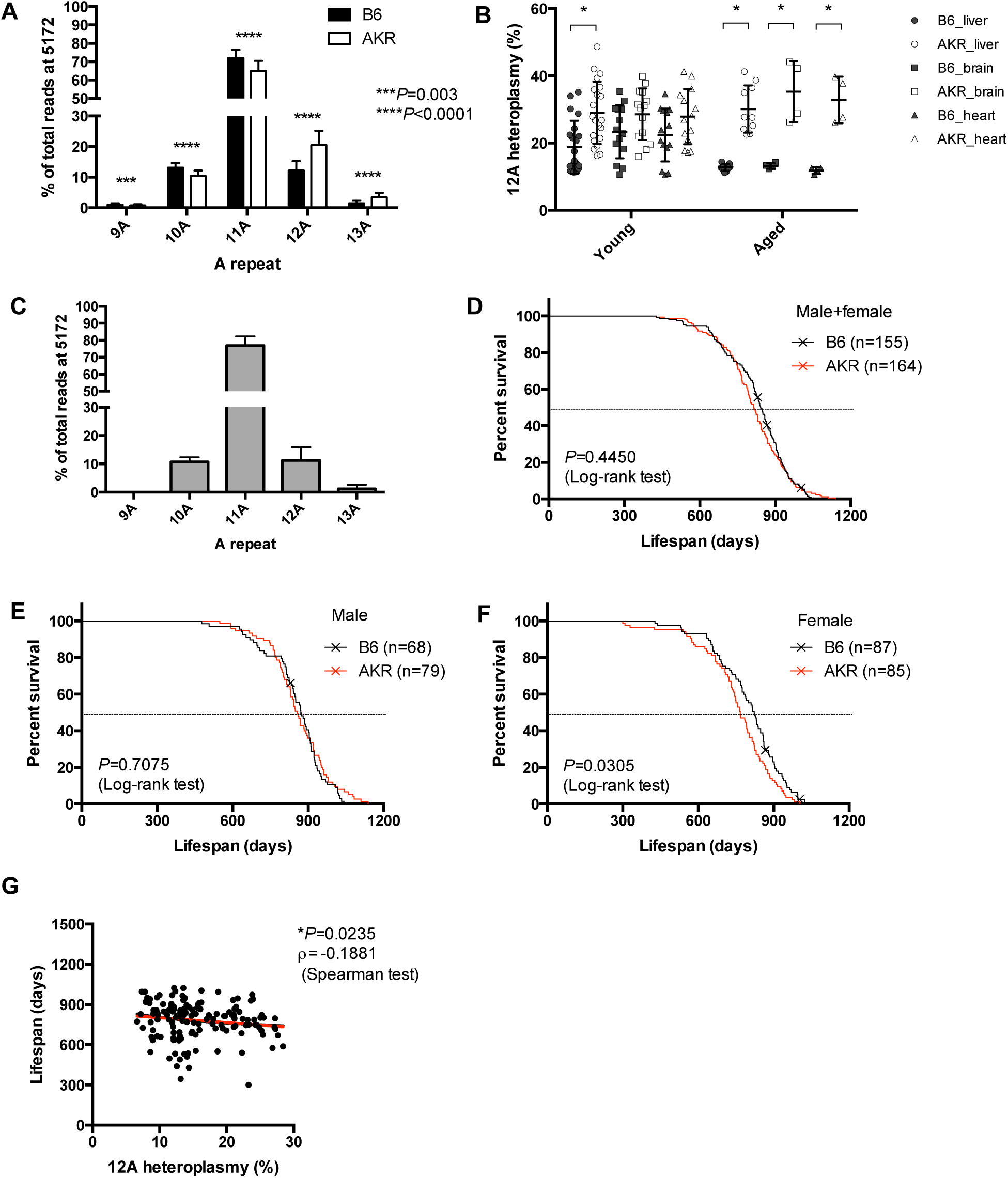
Levels of 12A heteroplasmy at 5172 in the origin of light strand replication negatively correlate to lifespan in mice. **(A)** Next generation sequencing of blood DNA obtained from moribund B6 and B6-mt^AKR^ (AKR) mice shows that AKR exhibits significantly higher levels of the 12A heteroplasmy at 5172 in the origin of light strand replication (OriL) compared with B6 (\*\*\**P=*0.003, \*\*\*\**P<*0.0001; two-way ANOVA). **(B)** The higher levels of 12A heteroplasmy in OriL are observed in different tissues (liver, brain and heart). The difference becomes more prominent when mice are aged (17-22 months old). \**P<*0.05, multiple *t-*test. **(C)** The heteroplasmy at 5172 is also detected in wild house mice (*Mus musculus domesticus*) caught on farms (n=215). **(D-F).** Survival curve of B6 and B6-mt^AKR^ (AKR) in both sexes (**d**), males (**e**), and females (**f**). Female AKR display a significantly shorter lifespan compared with B6 (*P=0.0305*, log-rank test). **(G)** The higher 12A heteroplasmy levels in OriL correlate with the shorter lifespan (ρ= -0.1881 *P=*0.0235, Spearman test).

### Levels of 12A heteroplasmy at 5172 in the OriL negatively correlate with lifespan

To investigate the impact of heteroplasmy at the OriL on murine lifespan and aging-related phenotypes, we conducted a longevity study using a large cohort of this strain, and B6 control mice. Female B6-mt^AKR^ live significantly shorter than B6 mice (approx. 50 days, *P=0.0305*, log-rank test), while the lifespan of males does not differ (**Figures 1D to F**, and **Tables S1A to D, Figure S1A to C**). Neither the aging score (i.e. presence of alopecia, greying hair, bod mass reduction, and kyphosis, as previously described (Ross et al., 2013)), tumor incidence, nor incidence of ulcerative dermatitis is significantly different between the strains (**Figures S1D to F)**. Sequencing of genomic DNA derived from blood obtained from moribund female mice (B6; n=84, B6-mt^AKR^; n=61) reveals an inverse correlation between lifespan and higher levels of heteroplasmy (**Figure 1G**, *ρ*=-0.1881, *P*=0.0235, Spearman test).

### Higher 12A heteroplasmy at 5172 in the OriL reduces the mtDNA/nDNA ratio, while the expression level of the mtDNA-encoded genes is increased

Next, we aimed to determine the functional consequence(s) of 12A heteroplasmy. Since the OriL is essential for mtDNA replication (Wanrooij et al., 2012), the ratio of mtDNA to nuclear genome copies was investigated by quantifying *mt-Co1* and *Vdac1* copies, respectively. We observe an inverse correlation with levels of 12A heteroplasmy (*ρ*=-0.4264, *P=*0.0025, Spearman test, **Figure 2A**), while the overall ratio is unaltered between B6-mt^AKR^ and B6 at different ages (**Figure 2B**). This is likely due to the high variability of 12A heteroplasmy in individual mice within each strain. This 12A-correlated phenomenon is also observed for another mitochondrially-encoded gene, *mt-Nd5*, again normalized to nuclear *Vdac1* (**Figures 2C and D**). Thus, higher levels of 12A heteroplasmy at 5172 correlate with lower mtDNA copy number. On the other hand, the expression of mitochondrial *mt-Co1* positively correlates with 12A heteroplasmy (*ρ=*0.5646, *P=*0.0033, Spearman test, **Figure 2E**), while no significant difference is observed between strains (**Figure 2F**). Consistent with the latter observation, the level of several OXPHOS proteins as well as the nuclear-encoded beta-actin protein is similar between strains (**Figures 2G and H**). These findings suggest that while higher levels of 12A heteroplasmy in the OriL lead to a reduction of mtDNA copy number, the cells are capable of a compensatory increase in expression of genes encoded by mtDNA despite its decreased copy number.

**Figure 2:**
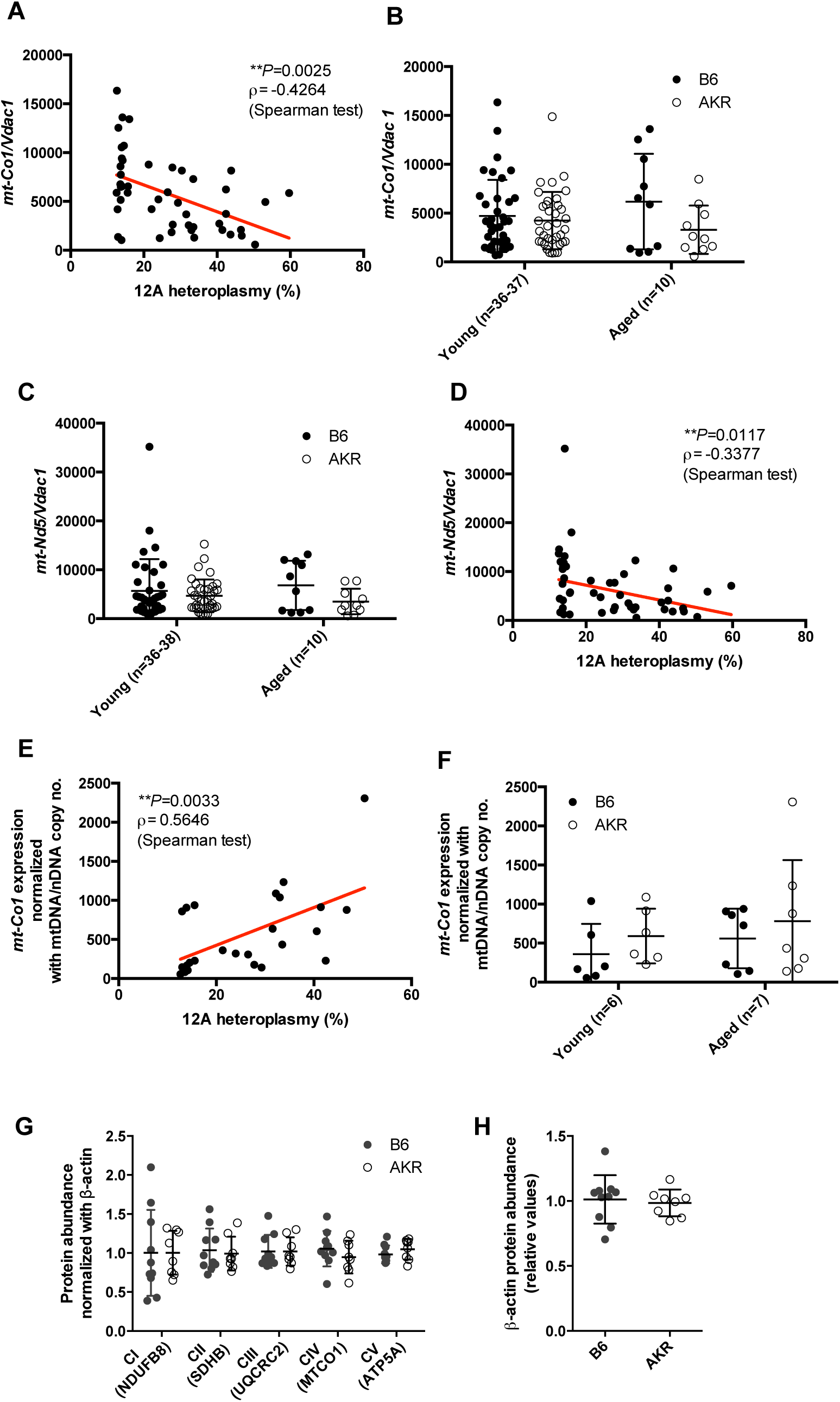
Higher levels of 12A heteroplasmy at 5172 in the origin of light strand replication reduces mtDNA/nDNA copy number ratio, while the expression levels of the mtDNA-encoded gene are increased. **(A)** The ratio of mtDNA (*mt-Co1*)- to nDNA (*Vdac1*) copy number was determined using liver genomic DNA obtained from B6 and B6-mt^AKR^ (AKR) mice (females, young: 3 months of age, aged: 18-22 months of age). The copy number ratio (*mt-Co1/Vdac1*) negatively correlates to levels of 12A heteroplasmy in OriL. N=48, *ρ=* -0.4264, \**P=*0.0025, Spearman test. **(B)** The values presented in **a**. were compared between strains, for which no difference is observed between B6 and AKR in each age group. **(C)** MtDNA gene copy number ratio based on *mt-Nd5/Vdac1* shows the same trend found by *mt-Co1/Vdac1*. **(D)** Higher 12A heteroplasmy levels in OriL also correlate with the copy number ratio of *mt-Nd5/Vdac1*. **(E)** The expression levels of *mt-Co1* in liver mitochondrial RNA were quantified using droplet digital PCR. The *mt-Co1* expression normalized to the *mt-Co1* copy number ratio reveals a significant correlation with the levels of 12A heteroplasmy. ρ*=*0.5646, \*\**P=*0.0033, Spearman test. **(F)** The values presented in **e**. display no significant difference when compared between the strains in each age group. **(G)** Quantified values of Western blotting of liver samples reveal unaltered protein levels of mitochondrial OXPHOS subunits. Females, three months of age, n=10 (B6), n=8 (AKR). **(H)** The same samples tested in **g**. were quantified for beta-actin.

### Mitochondrial function in young mice correlates to 12A heteroplasmy in the OriL when normalized to mtDNA copy number

Mitochondrial functional analysis including OXPHOS complex activity and hydrogen peroxide measurement assays display no significant correlation with 12A heteroplasmy in the OriL (**Figures S2A and B**). However, when normalized to the mtDNA to nDNA copy number ratio, 12A heteroplasmy significantly correlates with mitochondrial OXPHOS complex I, IV and V activity (complex I ρ=0.6214, *P*=0.0134; complex IV ρ=0.7321, *P*=0.0019; and complex V ρ=0.5668, *P*=0.0276, Spearman test, **Figure 3A**) as well as ROS levels (ρ=0.575, *P*=0.0249, Spearman test, **Figure 3B**). This phenomenon is observed in young mice only (three to six months of age; **Figures 3A and B**), while an aged population maintains the trend of this correlation (around 18-20 months of age; **Figure S3A and B**).

**Figure 3:**
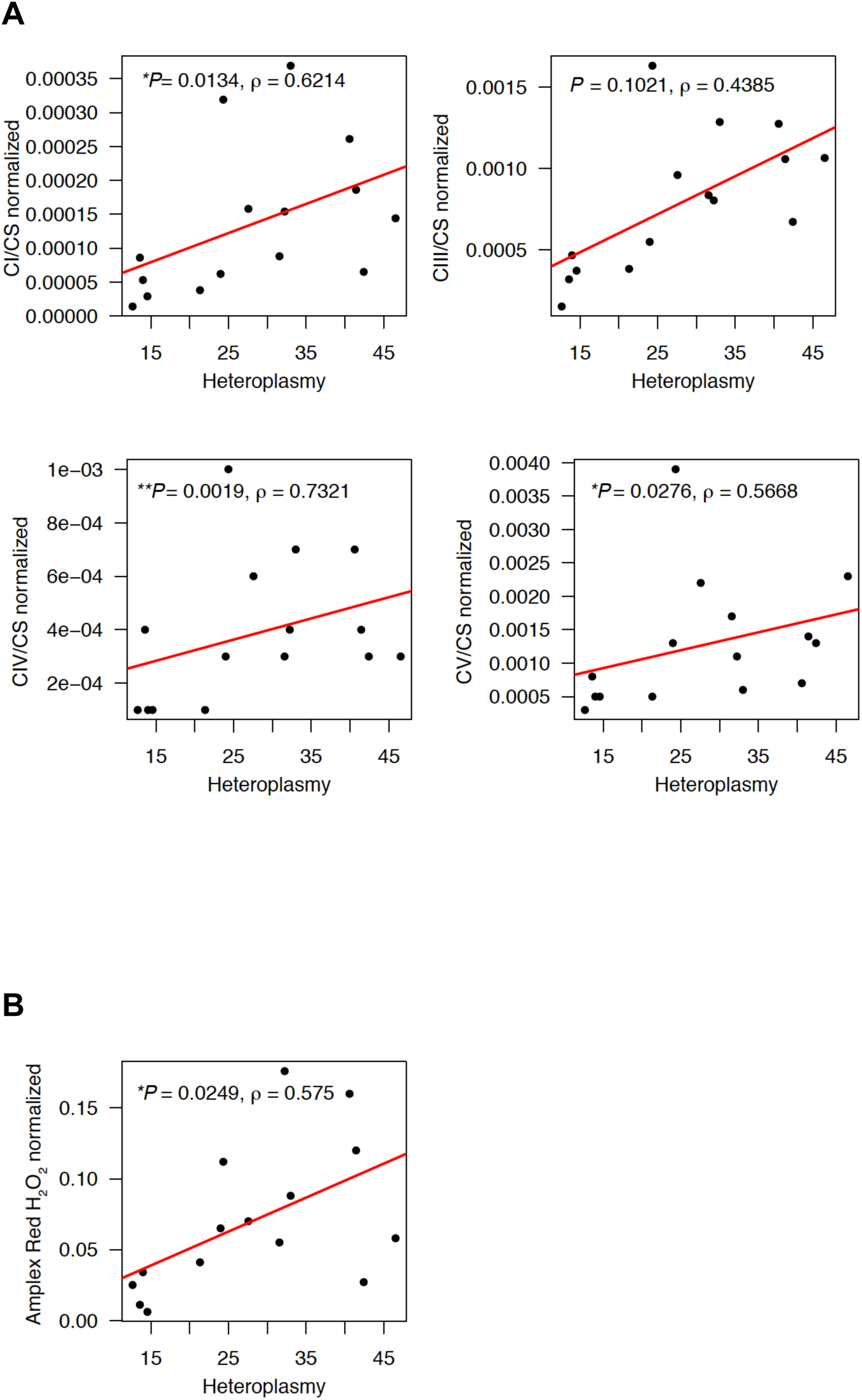
Mitochondrial functions in young mice correlate to the levels of 12A heteroplasmy at OriL. **(A)** Oxidative phosphorylation (OXPHOS) complex activities measured using liver mitochondria obtained from young mice (three months old, females), normalized to the individual mtDNA copy number ratio (*mt-Co1/Vdac1*). Complex I (CI) activities were normalized to citrate synthase (CS) activities. Complex III (CIII), complex IV (CIV) as well as complex V (CV) values were also analyzed in the same manner. Complex I (CI) activities normalized to citrate synthase (CS) activities significantly correlate with the levels of 12A heteroplasmy in OriL when normalized to the individual mtDNA copy number ratio. N=7 (B6), n=8 (B6-mt^AKR^). Correlations were investigated using the Spearman rank correlation. **(B)** Levels of hydrogen peroxide in female liver mitochondria obtained from young mice (3-5 months old) normalized to the individual mtDNA copy number ratio (*mt-Co1/Vdac1*). The normalized ROS levels show a positive correlation with the levels of 12A heteroplasmy in OriL (ρ=0.575, *P*=0.0249, Spearman test). N=7 (B6), n=8 (B6-mt^AKR^).

### Impaired glucose and lipid metabolism may determine healthspan in B6-mt^AKR^ mice

Mitochondrial metabolism has a well-defined impact on glucose metabolism (Maassen et al., 2004; Wiederkehr and Wollheim, 2006), and conversely, increased glucose levels promote systemic aging (Yin et al., 2016). We thus lastly evaluated the putative impact of 12A heteroplasmy in the OriL on the metabolic phenotype of both B6-mt^AKR^ and B6 mice. Random-fed plasma glucose levels are higher in 12A heteroplasmy B6-mt^AKR^ mice than B6 controls (**Figure 4A**). Consistently, plasma fructosamine concentrations, which reflect glucose levels over a 2-3 week period prior to measurement, are also increased in states of high 12A heteroplasmy (**Figure 4A**), indicating a long-term elevation of plasma-glucose in these mice. Moreover, B6-mt^AKR^ display a consistent elevated oral glucose tolerance test (**Figure 4B**). Nevertheless, insulin levels in both fasted- and random-fed plasma samples are unaltered, as are lactate, cholesterol and fasted glucose levels (**Figures S4A and B**). Further, comparable levels of glycogen storage contrasted by higher activity of pyruvate kinase (PK, indicative for glycolysis) and phosphoenolpyruvate carboxykinase (PEPCK, indicative for gluconeogenesis) (**Figure 4C**) indicate an increase in glucose consumption in B6-mt^AKR^ mice. Higher levels of free fatty acids are also present in B6-mt^AKR^ (**Figure 4C**), suggesting a global reduction of beta-oxidation in these mice. This is supported by our findings of elevated long-chain fatty acids in liver samples of B6-mt^AKR^ (**Figure 4D** and **Figure S4C**), consistent with an impaired capacity for mitochondrial metabolism, and in particular beta-oxidation. Thus, mice with higher 12A heteroplasmy (B6-mt^AKR^) exhibit skewed glucose and lipid metabolism, and that may be considered causal for the impaired lifespan of B6-mt^AKR^ mice in comparison to low-heteroplasmy B6 control mice. This is consistent with long-standing published evidence that mitochondrial mutations, as well as impaired mitochondrial function, impair glucose tolerance and can cause so-called mitochondrial diabetes mellitus, as reviewed elsewhere (Maassen et al., 2004; Wiederkehr and Wollheim, 2006).

**Figure 4:**
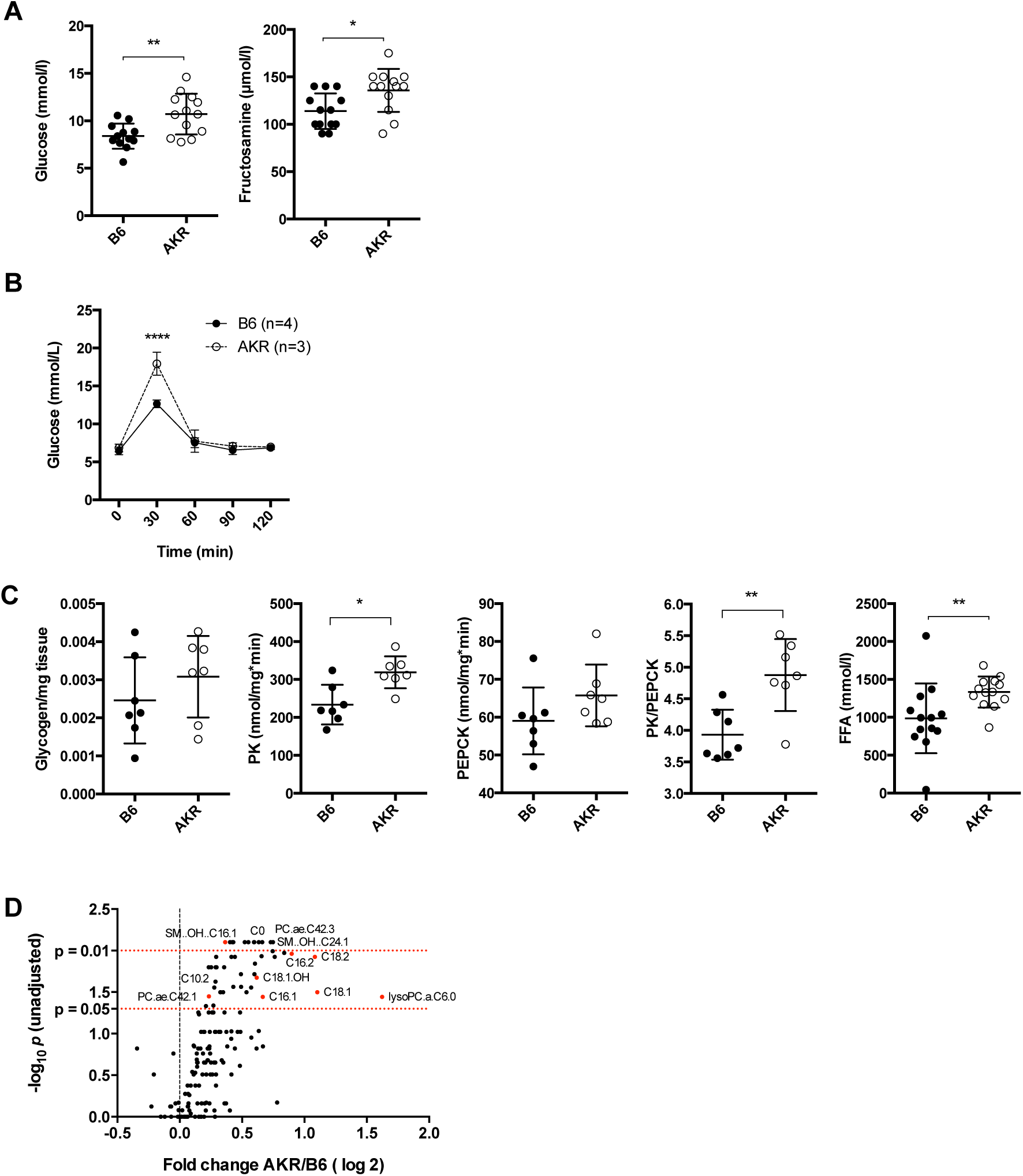
Impaired glucose and lipid metabolism in B6-mt^AKR^ mice. **(A)** Glucose and fructosamine levels were determined in plasma samples from random-fed B6 and B6-mt^AKR^ mice (3M, females). \**P=* 0.0106 (fructosamine), \*\**P=*0.0045 (glucose), Mann-Whitney U test. N=12-13 per strain. AKR=B6-mt^AKR^. **(B)** Oral glucose tolerance test (2g/kg, p.o.) was performed in young (3 months old) B6 and B6-mt^AKR^ female mice. *\*\*\*\*P<0.0001*, two-way ANOVA. **(C)** Glycogen content, activities of pyruvate kinase (PK) and those of phosphoenolpyruvic carboxykinase (PEPCK), and the ratio of PK to PEPCK were determined in liver samples obtained from the mice tested in **a**. Free fatty acids levels were measured in plasma samples obtained from overnight-fasted B6 and B6-mt^AKR^ (3M, females, same animals tested in **a**). \**P*=0.0111 (PK), \*\**P*=0.0041 (PK/PEPCK), \*\**P=*0.0027 (FFA), Mann-Whitney U test. **(D)** Liver lipid metabolites (free carnitine, acylcarnitines, phosphatidylcholines, lysophosphatidylcholines and sphingomyelines) were measured in B6 and B6-mt^AKR^. AKR=B6-mt^AKR^. Females, n=5/strain.

### The gene expression pattern in mice with high levels of 12A heteroplasmy at the OriL is similar to that in advanced age mice

Next, to elucidate the pathways involved in heteroplasmy-related phenotypes, RNA-seq was performed on liver tissue from 3-4 and 19-20-month-old B6 and B6-mt^AKR^ mice. Of 14,031 expressed genes, 103 are differentially expressed between young B6-mt^AKR^ and B6 mice (*p<0.01*, **Figure 5A, Data S1**). Genes highly expressed in mice with higher heteroplasmy include *Mtor* and *Fbxo32*, suggesting that heteroplasmy exhibits a mild, but significant impact on the mTOR pathway. Interestingly, *Fbxo32* (Atrogin-1; Mafbx) was shown to be upregulated in sarcopenia. Pathways leading to sarcopenia include a down regulation of the PI3K/AKT pathway and activation of the FOXO transcription factor (Sandri et al., 2004).

**Figure 5:**
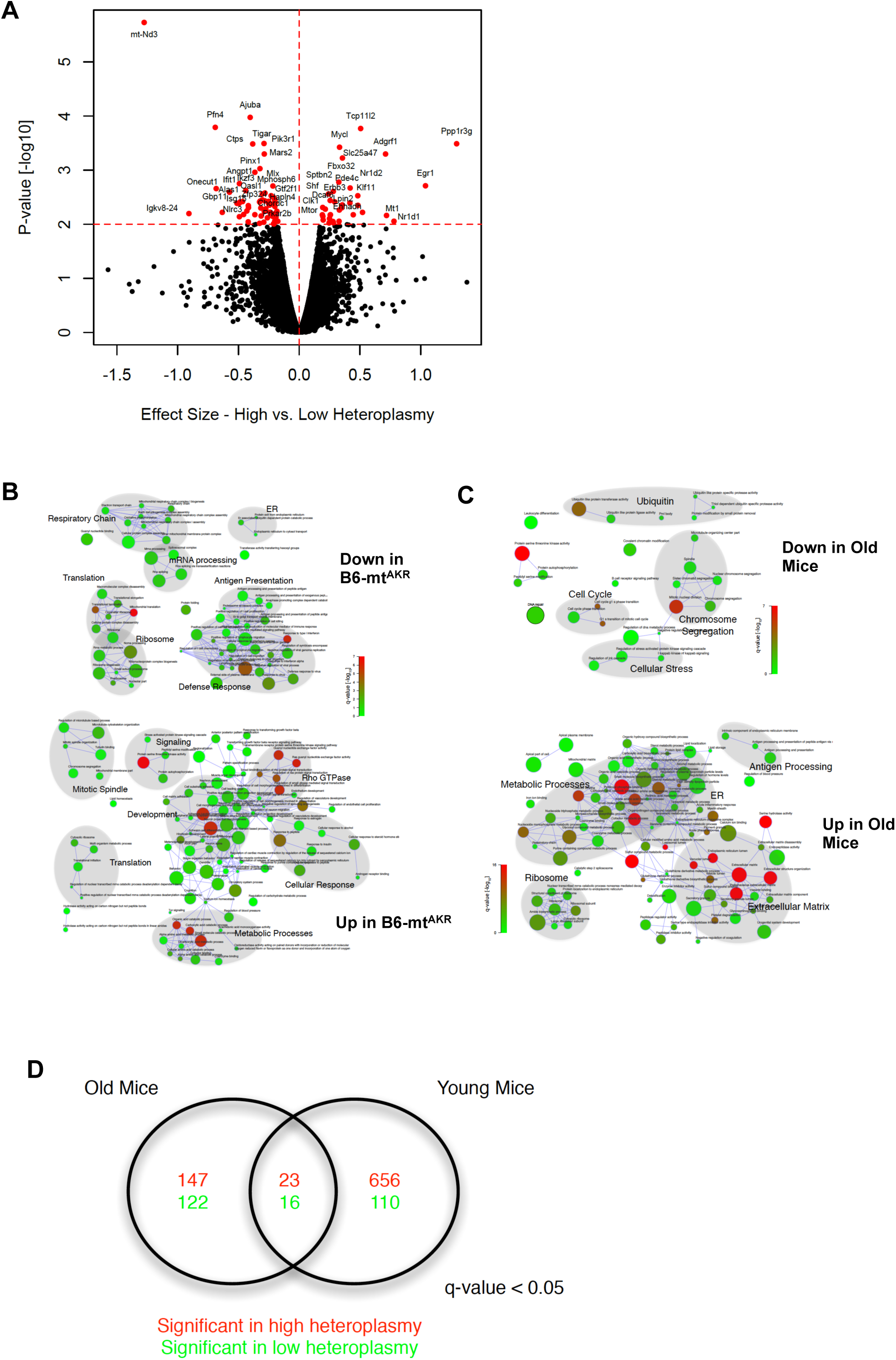
RNA-seq analysis of liver samples of B6-mt^AKR^ and B6 displayed differential gene expression between the strains. **(A) Differentially expressed genes between B6-mt^AKR^ and B6 mice.** The volcano plot demonstrates the effect size versus the –log10 *p-*value of differentially regulated genes. Genes whose expression were *p<0.01* were plotted in red. All young and aged mice were included in the analysis. **(B) Gene set enrichment analysis of genes whose expression was affected by the 12A heteroplasmy levels.** The network represents the significant gene sets as nodes; with the node color corresponding to the significance and the node diameter to the gene set size. Nodes are connected by edges if they share at least 20% of their genes. Areas highlighted in gray background indicate the most prominent function in the clustered nodes. **(C) Gene set enrichment analysis of genes between young and aged mice.** The same method as (**B**) was used for data analysis. **(D) Biological process ontology terms commonly affected by aging and 12A heteroplasmy levels.** Of 23 significantly up-regulated biological processes controlled by both aging and heteroplasmy, metabolic processes were up-regulated in aged mice and mice with higher 12A heteroplasmy levels. On the other hand, the mitochondrial processes were down-regulated.

Gene set enrichment analysis (GSEA) reveals that up-regulated genes include those involved in metabolic processes and cellular responses (mainly to hormones), while down-regulated genes include those involved in the respiratory chain, ribosomes and translation (e.g. mitochondrial translation). (**Data S2**, **Figure 5B**). We also evaluated whether aging and heteroplasmy play a complementary role in driving gene expression. The overlap between GSEA according to heteroplasmy versus age (**Figure 5C**) results in 23 common highly expressed gene ontology (GO) terms and 16 down-regulated terms (**Figure 5D**). Glucose metabolism and cell adhesion were prevalent in higher heteroplasmy (B6-mt^AKR^) and aged mice, while mitochondrial processes were prevalent in lower heteroplasmy (B6) and young mice (**Data S3**). These transcriptomic data are in agreement with the findings of mtDNA copy number and gene expression (i.e. less mtDNA copy number and higher mtDNA-gene expression in B6-mt^AKR^; **Figure 2A, D and E**), as well as the metabolic phenotype data, i.e. genes involved in metabolic processes are up-regulated, presumably in order to control glucose metabolism in mice with higher levels of 12A heteroplasmy (**Figures 4 and Figure S4**).

### The impact of low-level heteroplasmic mutations depends on the location

Lastly, we evaluated whether another heteroplasmic mutation present in the mice has an impact on lifespan. The sequencing data revealed that both B6 and B6-mtAKR strain carry a heteroplasmic mutation at nt9821, in the tRNA-arginine gene (*mt-Tr*). Almost all B6 samples are 8-adenine repeat (8A) homoplasmic, while 9A is the major genotype in B6-mt^AKR^ with a very low copy number of 8A (**Figure S5A**). However, the level of heteroplasmy in *mt-Tr* does not correlate to lifespan in our study (**Figure S5B**), indicating that the impact of lower heteroplasmy on lifespan is highly dependent on the location.

## Discussion

Individual mammalian cells, including oocytes, carry thousands of individual copies of mitochondrial DNA. These copies may differ in regards to individual bases or entire regions. This so-called heteroplasmy of mitochondrial DNA has been described many years ago (Solignac et al., 1983; Hauswirth et al., 1984; Boursot et al., 1987) and is known to cause mitochondrial dysfunction if affecting a high percentage of mtDNA copies, i.e. over 70-80% (Larsson, 2010; Stewart and Chinnery, 2015). Here, we observed “low-level” heteroplasmy at nt5172 in the OriL (>20%) in the B6-mt^AKR^ mouse line, which is more pronounced than the degree of heteroplasmy observed in the B6 background (namely ∼ 10%).

This is the first report demonstrating the impact of naturally occuring low-level heteroplasmic mutations, and specifically in the OriL locus, on mammalian aging. The OriL is indispensable for mtDNA maintenance in mice (Wanrooij et al., 2012). The OriL sequence forms a stable stem-loop structure when mtDNA is single-stranded during replication, and mtDNA-directed RNA polymerase (POLRMT) initiates primer synthesis from the polyA stretch of the loop (Fusté et al., 2010). Since the region is critical for mtDNA replication and maintenance, the OriL appears to be well protected from the incidence of mutations (Wanrooij et al., 2012). Our transcriptome data demonstrates that the lower the level of 12A heteroplasmy in the OriL, the higher the up-regulation of mitochondrial translation (**Figure 5B**), supporting a functional link between the level of 12A heteroplasmy in the OriL and mtDNA copy number (**Figure 2A and D**). Additionally, the particular region seems to be a specific target of the nematodal CLK-1 protein (Gorbunova and Seluanov, 2002). The same study demonstrates that its mouse homologue also displays binding activity specific to the OriL loop, suggesting a regulatory function of CLK-1 in mtDNA replication in addition to its known ubiquinone biosynthesis function (Gorbunova and Seluanov, 2002). Mutations in the *clk-1* gene result in an extension of lifespan in nematodes (Lakowski and Hekimi, 1996), and a partial inactivation of the mouse homologue *Mclk-1* also extends lifespan in mice (Liu et al., 2005). Global deletion of *Mclk-1* in adult mice results in decreased levels of fasting glucose and triglycerides (Wang et al., 2015), suggesting the potential role of the OriL in metabolic control. This experimental evidence supports our observation regarding the correlation between heteroplasmic mutations in OriL and lifespan in mice.

An additional low-level heteroplasmy at position 9820 is present in our data, in both B6 and B6-mt^AKR^ mice. This position is located in the tRNA-arginine gene (*mt-Tr*), which is known to be polymorphic in common inbred strains (Bayona-Bafaluy et al., 2003; Johnson et al., 2001; Sachadyn et al., 2008). No correlation is observed between the heteroplasmic mutation in *mt-T*r and any of the phenotypes we investigated, thus supporting our conclusion that the impact of low-level heteroplasmy is dependent on the location within the mtDNA, OriL in this study.

In summary, we present experimental evidence linking the degree of mtDNA heteroplasmy to murine lifespan. Specifically, higher levels of 12A heteroplasmy at 5172 in the OriL correlate with shorter lifespan and impaired glucose metabolism in mice. Furthermore, this heteroplasmy influences mtDNA copy number. In young mice, such a reduction of mtDNA copy number can be compensated for by increased expression of the affected genes. In contrast, this compensatory mechanism is missing in aged mice, which tend to carry increased levels of 12A heteroplasmy. The heteroplasmic mutation investigated in this study is natural, low-level and stable over generations. We demonstrate that such quiescent variation in mtDNA influences healthspan even under normal housing condition, suggesting that additional environmental triggers such as metabolic stress could amplify the phenotypic consequences. Further studies to elucidate the involved mechanisms, particularly the interaction between mtDNA variants and environmental stress factors, are warranted. Given the established role of mtDNA variations in human exercise performance, the current findings may likely translate into a co-determining function of human aging.

## Acknowledgements

The authors thank Miriam Daumann, Miriam Freitag, Beate Laube, Doris Pöhlmann, Melanie Vollstedt, and Nicole Thomsen for excellent technical assistance. We thank Stephanie Derer and Christian Sina for valuable discussion. This work was supported by grants from the Bundesministerium für Bildung und Forschung (BMBF, 0315892B), DFG (EXC 306 and CRC 1182), University of Lübeck (P01-2012), the Swiss National Science Foundation (Schweizerischer Nationalfonds, SNF 31003A_156031), and the European Union´s Horizon 2020 research and innovation program (633589).

## Author Contributions

M.H. and S.M.I. designed the study. M.H., P.S. and J.Y. performed the lifespan study, mitochondrial functional study as well as mitochondrial genome sequencing experiments and analyzed the data. M.N.W. and A.Z. conducted the statistical analysis of the survival data together with A.K. and H.B.; Y.G. analyzed the mitochondrial sequencing data. R.H. performed RNA-seq. A.K. and H.B. analyzed transcriptome data together with G.F.; M.V., M.B., and J.F.B. collected wild mice samples and analyzed the data with D.T.; K.Z., K.J., R.O., and C.P. performed metabolic phenotyping experiment and analyzed the data. J.M., A.O., O.J., H.L., M.S., M.R., J.A., and R.K. contributed to interpretation of the metabolic phenotype data. K.S. and J.R. contributed to interpretation of the mitochondrial phenotype data. M.H., J.F.B., M.R. and S.M.I. wrote the manuscript with contributions from all other authors. S.M.I. directed the study.

